# 7-ketocholesterol contributes to microglia-driven increases in astrocyte reactive oxygen species in Alzheimer’s disease

**DOI:** 10.1101/2025.01.19.633810

**Authors:** Kayalvizhi Radhakrishnan, Yiyu Zhang, Oluwaseun Mustapha, Thaddeus K. Weigel, Clint M. Upchurch, Xiaodong Tian, Franklin Herbert, Wenyuan Huang, Norbert Leitinger, Ukpong B. Eyo, Huiwang Ai, Heather A. Ferris

**Author notes:** Co-corresponding authors: Correspondence to: Heather A. Ferris, MD, PhD, Division of Endocrinology and Metabolism, University of Virginia, PO Box 801406, Charlottesville, VA, 22908, USA., Huiwang Ai, PhD, Department of Molecular Physiology and Biological Physics, University of Virginia, PO Box 800736, Charlottesville, VA, USA. Authors contributed equally (Kayalvizhi Radhakrishnan, Yiyu Zhang).

## Abstract

Oxidative stress is a prominent feature of Alzheimer’s disease. Within this context, cholesterol undergoes oxidation, producing the pro-inflammatory product 7-ketocholesterol (7-KC). In this study, we observe elevated levels of 7-KC in the brains of the 3xTg mouse model of AD. To further understand the contribution of 7-KC on the oxidative environment, we developed a method to express a genetically encoded fluorescent hydrogen peroxide (H_2_O_2_) sensor in astrocytes, the primary source of cholesterol in the brain. With this sensor, we discovered that 7-KC increases H_2_O_2_ levels in astrocytes in vivo, but not when directly applied to astrocytes in vitro. Interestingly, when 7-KC was applied to a microglia cell line alone or mixed astrocyte and microglia cultures, it resulted in microglia activation and increased oxidative stress in astrocytes. Depletion of microglia from 3xTg mice resulted in reduced 7-KC in the brains of these mice. Taken together, these findings suggest that 7-KC, acting through microglia, contributes to increased astrocyte oxidative stress in AD. This study sheds light on the complex interplay between cholesterol oxidation, microglia activation, and astrocyte oxidative stress in the pathogenesis of AD.

## Introduction

Astrocytes are the main source of cholesterol in the brain (Ferris et al., 2017). Astrocyte-derived cholesterol can be used in cell membranes and organelles of the astrocyte, stored in lipid droplets, or transported to other cells of the brain. A variety of metabolic stressors, including excess lipid accumulation (Copetti-Santos et al., 2015), amyloid beta (Aβ) exposure (González-Reyes et al., 2017) and cytokine exposure (Sheng et al., 2013), can result in increased reactive oxygen species (ROS) in astrocytes. 7-ketocholesterol (7-KC) is a cholesterol derivative produced as a result of non-enzymatic ROS-induced oxidation of the cholesterol molecule at the C-7 position. Elevated levels of 7-KC have been detected in atherosclerotic plaques, Alzheimer’s disease (AD) brains, the serum of individuals with diabetes (Pathak et al., 2024), and the retina of individuals with age-related macular degeneration (Indaram et al., 2015). 7-KC is known to trigger inflammatory pathways and induce apoptosis in neurons and glial cells (Nury et al., 2021; Testa et al., 2016; Yammine et al., 2020; Zarrouk et al., 2021). As a toxic component, 7-KC further promotes oxidative stress, resulting in an amplification of ROS production, which can damage proteins, lipids, and DNA. While there are several studies of 7-KC in microglia, the impact on astrocytes has not yet been assessed.

Direct measurement of ROS in biological systems presents significant challenges. The most common techniques involve the quantification of oxidized lipids or proteins, the end products of oxidative stress. These measures cannot dynamically assess ROS levels, instead providing a snapshot of accumulated exposure.

Furthermore, it is often challenging to gain spatial and cell-specific information from the analyzed samples. Recently, a group of genetically encoded fluorescent redox indicators (GERIs) has been developed for the spatial and temporal monitoring of ROS (Pak et al., 2020; Pang et al., 2021). However, they have not been widely utilized to study disease models, particularly AD models. In our previous study, we fused the mitochondrial targeting sequence to a ratiometric hydrogen peroxide sensor, HyPerGR_T_ (Li et al., 2022). HyPerGR_T_ is comprised of a green fluorescent hydrogen peroxide sensor, HyPer7 (Pak et al., 2020), and a red fluorescent protein, tdTomato (Shaner et al., 2004) (Pak et al., 2020). This fusion enabled ratiometric imaging of mitochondrial hydrogen peroxide in cultured HeLa cells, primary mouse neurons, Aβ-treated organotypic brain slices, and brain slices from Aβ-treated mice (Li et al., 2022).

In this study, we expanded the use of HyPerGR_T_ and developed an adeno-associated viral (AAV) vector to specifically express the sensor in the cytoplasm of astrocytes for specific detection of astrocyte ROS. We coupled this new tool with other techniques, including mass spectrometry, *in vitro* astrocyte and microglial culture, fluorescence imaging, and AD mouse models to study the roles of 7-KC in astrocytes in the pathogenesis of AD.

We identified that both astrocyte ROS and 7-KC are elevated in the brains of the 3xTg mouse model of AD. We further show that 7-KC, but not cholesterol, increases astrocyte hydrogen peroxide levels *in vitro* and *in vivo*. To determine if the elevation in astrocyte ROS was a direct effect of 7-KC on astrocytes, we applied 7-KC to primary astrocyte cultures *in vitro*. Our findings revealed a limited direct impact of 7-KC on astrocytes. In contrast, 7-KC was able to activate microglia *in vitro*. Furthermore, when mixed glial primary cultures were treated with 7-KC, there was astrocyte activation and an increase of astrocyte ROS. Finally, depletion of microglia from 3xTg mice reduced 7-KC in the cortex of these animals. Overall, this study shows that 7-KC is increased in the AD brain and can contribute to increased astrocyte ROS by activating microglia. In addition, this study introduces an astrocyte-specific cytosolic AAV redox indicator as a convenient tool for studying ROS specifically in the cytosol of astrocytes both *in vitro* and *in vivo*.

## Materials and Methods

### General animal information

Animal experiments were performed in compliance and with the approval of the University of Virginia Institutional Animal Care and Use Committee. 3xTg experiments were conducted with 8–14 month-old-females 3xTg (Oddo et al., 2003) (B6;129-*Psen1^tm1Mpm^* Tg(APPSwe,tauP301L)1Lfa/Mmjax, Jackson Laboratory #034830), and age-matched B6129SF2/J (Jackson Laboratory #101045) females as wild-type controls. Experiments with 5xFAD mice (B6.Cg-Tg(APPSwFILon,PSEN1*M146L*L286V)6799Vas/Mmjax, Jackson Laboratory # 034848) were performed on 8-9 month old heterozygous males and non-transgenic male littermates were the controls. C57BL/6J mice were obtained from Jackson Laboratory (#000664) and maintained as pure colonies.

### Depletion of microglia in animals

For microglia depletion experiments, 14-month-old female 3xTg and B61292F2/J wild-type mice were given chow formulated with PLX3397 (660 mg/kg) or control chow. They were maintained on treatment or control diets for four weeks.

### Measurement of 7-KC levels by mass spectrometry

For 7-KC measurements, mice were first anesthetized with a ketamine-xylazine solution and transcardially perfused with chilled phosphate-buffered saline (PBS). All mice were sacrificed from Zeitgeber time 5–9. The brain region of interest was dissected out, weighed, and flash-frozen. Samples were homogenized in water with 1.2% butylated hydroxytoluene (BHT) (Sigma-Aldrich) to prevent oxidation. Sterols were extracted using the modified Bligh-Dyer method. Samples were mixed with deuterated standards (cholesterol-d7 and 24-dehydro cholesterol-d6, Cayman Chemical Company) to control for differences in extraction efficiency. Samples were further extracted with chloroform, dried under nitrogen gas, and resuspended in methanol. Sterols were separated using binary reverse-phase ultra-high performance Vanquish liquid chromatography (Phenomenex Reverse Phase C18, 3 mm particle size, 2 mm diameter, 250 mm length) using Solvent A (methanol; 5 mM ammonium acetate) and Solvent B (15% water; 85% methanol; 5 mM ammonium acetate) with a flow rate of 0.200 mL/min and the following gradient: 0 to 2 minute, 0% B; 2 to 8 minutes, 0-100% B; 8-32 minutes, 100% B, 32 to 35 minutes, 100-0% B; 35 to 40 minutes 0% B. Column temperature of 35°C was used, and sterols were quantified by electrospray ionization (ESI) spectrometry and selective ion monitoring (SIM) using a Thermo Fisher Q Exactive mass spectrometer. The mass spectrometer was operated with the following settings: Resolution – 35,000; AGC Target – 5e5; Scan Range – 150-2000 m/z; Isolation window – 2.0 m/z; and Runtime – 10 to 22 minutes.

### Mouse primary cultures

Astrocyte-enriched mixed glial cultures were prepared from cortices of P1 to P3 C57BL/6J mouse pups. The brains were dissected in ice-cold Hanks’ Balanced Salt Solution (HBSS) and the cortices were separated from other tissues with meninges removed completely.

Isolated cortices were dissociated with trypsin and DNase in a 37°C water bath for 30 min. The enzyme reaction was stopped by adding fetal bovine serum (FBS)-containing media to the dissociated tissue. The contents were centrifuged, and the cell pellet was resuspended and plated onto T75 flasks with DMEM containing 10% heat-inactivated FBS, 100 IU/mL penicillin, 100 μg/mL streptomycin, and 1.25 mM L-glutamine. After 7 days, the flask was shaken in an orbital shaker at 37°C at 500 RPM for 4 to 5 hours to remove microglia. The astrocytes were detached using trypsin and seeded onto 6-well plates for qPCR or 0.1 mg/ml poly-D-lysine coated glass chambers for imaging. To deplete microglia in designated experimental groups, 5 μM PLX-3397 in DMSO (Cat. # HY-16749, MedChem Express) was added at the time of cell seeding and replenished with each media change. For 7-KC treatments, 10 μM 7-KC or cholesterol dissolved in ethanol prepared as a 1000x stock was added to the culture media, the final concentration of ethanol in media was around 0.1%. 24 hours after the treatment, the cells were harvested for RNA extraction.

### In vitro imaging of astrocytes

Astrocytes were transduced with the AAV-HyPerGR_T_ and allowed one week for viral expression. The transduced cells were treated with 100 mM hydrogen peroxide or PBS control for 5 mins. In case of 7-KC, 10 μM 7-KC or cholesterol were applied to the cells for 24 hours. The cells were then washed three times with PBS and blocked with 4% paraformaldehyde containing 20mM N-Ethylmaleimide followed by three PBS washes. The cells were then mounted using Invitrogen’s Prolong Gold mounting media. The images were taken using Keyence slide scanner with standard wavelengths 488 nm for green and 561 nm for red channels. The green and red intensities of individual cells were measured using Fiji / ImageJ and the green to red ratio was calculated. As many as 100 cells from several fields taken from 3 different cultures for each condition were used for quantification.

### Culture and analysis of SIM-A9 microglia cells

SIM-A9 is a microglia-derived cell line (Nagamoto-Combs et al., 2014). SIM-A9 cells were cultured in DMEM-F12 containing 5% horse serum, 10% fetal bovine serum, 100 IU/mL penicillin, and 100 μg/mL streptomycin. Cells were primed with lipopolysaccharide (LPS) as previously described (Indaram et al., 2015). Briefly, the SIM-A9 cells were treated with 0.5 μg/ml LPS for 16 hours, followed by 12 μM 7-KC (Cayman Chemicals, Cat # 16339) or 12 μM cholesterol (Cayman Chemicals, Cat # 9003100) for 6 hours. At the end of the experiments, cells were harvested for RNA extraction and qPCR as described above.

### Quantitative PCR

Total RNA was extracted from cells using RNeasy Plus Mini Kit (Qiagen) followed by cDNA synthesis using an iScript cDNA Synthesis Kit (BioRad). qPCR was performed using iTaq Universal SYBR Green Supermix (BioRad). The primer sequences for qPCR are listed in Table 1. The mRNA expression levels were normalized to the housekeeping gene TATA binding protein. The relative gene expression was calculated using 2^(-delta Cq)^ and fold change was calculated using 2^(-delta delta Cq)^.

**Table 1.**
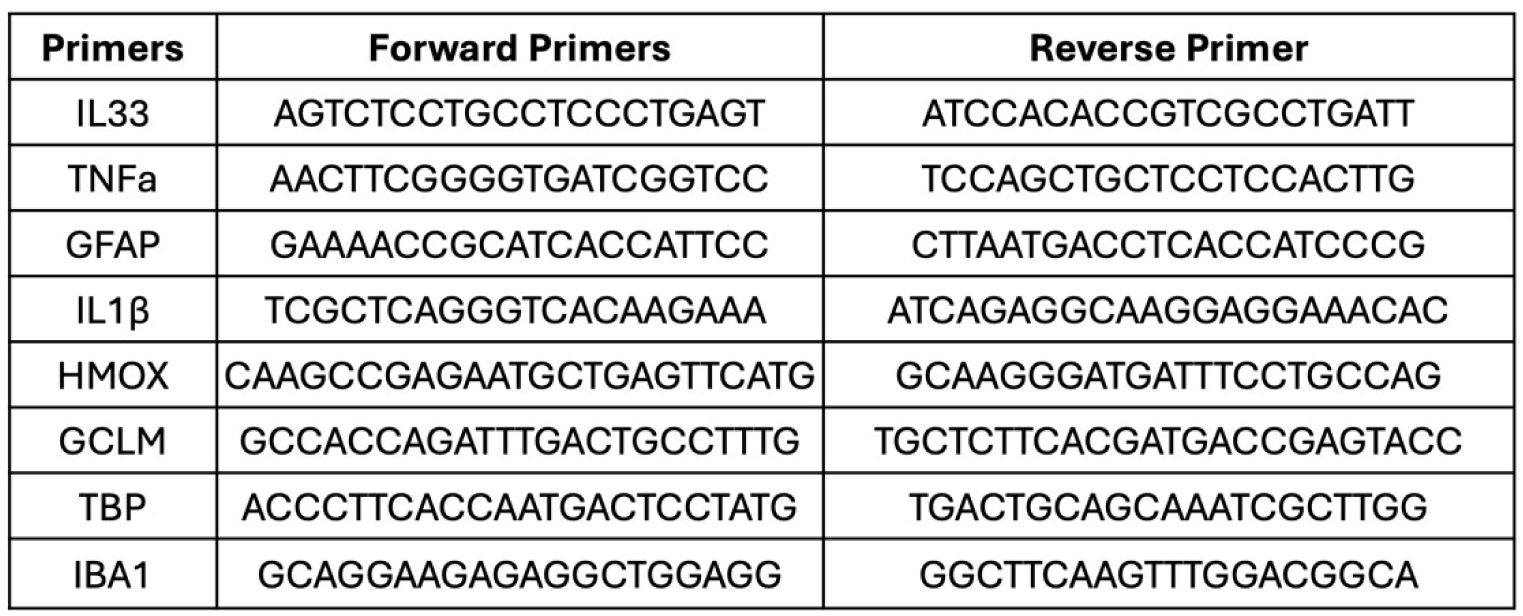

### Plasmid construction and viral packing

The construction of pAAV-hSyn-HyPerGR_T_ was described previously (Li et al., 2022). To use it for AAV packing and achieve astrocyte-specific expression, a truncated version of the glial fibrillary acidic protein (GFAP) promoter (GfaABC_1_D) was amplified from pTYF-1xGfaABC1D-EGFP (Addgene plasmid #19974) (Liu et al., 2008) and used to replace the hSyn promoter in pAAV-hSyn-HyPerGR_T_, resulting in a new plasmid which we named pAAV-GFAP-HyPerGR_T_. This plasmid was combined with pDeltaF6 (Addgene plasmid #112867) and pAAV2/5 (Addgene plasmid #104964) for viral packing and purification, following a described protocol (Tian et al., 2022). The viral titers were assessed using a qPCR method provided by Addgene. The AAV titers employed in this study were approximately 2 × 10^12^ genome counts (GC) per mL.

### Hippocampal injection

The AAV virus was administered to mice via stereotaxic injection. Briefly, animals were anesthetized using isoflurane in an induction chamber, before being transferred to a Stoelting stereotaxic apparatus where there was a continuous flow of isoflurane. 3 mg/kg of ketoprofen was given subcutaneously. The surgical site was shaved, disinfected with betadine and ethanol, and then infiltrated with buprenorphine (2.4 mg/kg). Craniotomies were done using a dental drill at the stereotaxic coordinates appropriate for the hippocampus; AP, −1.7 mm; ML, ±1.2 mm; DV, −1.5 mm relative to bregma (Tetteh et al., 2019). 1 μl of the virus was injected at a rate of 0.1 μL/min. After withdrawing the needle, the skin was sealed with tissue adhesive. Injected animals were monitored and given ketoprofen for five subsequent days for pain relief. In experiments with 7-KC injections, 1 μl of 5 μM of 7-KC or cholesterol or 1 μl of methyl-beta-cyclodextrin (MBCD) was injected as described above.

### Brain slice imaging and quantification

Imaging of prepared acute brain slices was done on a Scientifica SliceScope Pro 1000 equipped with a Photometrics Prime 95B scientific CMOS camera, a CoolLED Enhanced pE-300^white^ LED light source, and a 40× (NA 0.8) water immersion objective lens. The green and red channels were imaged using an FITC filter cube (470/40 nm excitation and 525/50 nm emission) and a TRITC filter cube (545/25 nm excitation and 605/70 nm emission). To process the ratios, individual cells were circled as regions of interest (ROIs) in ImageJ, and their intensity was measured in both channels. The green/red intensity ratio was then computed. Approximately 100 cells from several acute brain slices for each animal were analyzed.

### Statistical quantification

For qPCR and *in vitro* fluorescent ratiometric quantification experiments two-tailed unpaired t-test was performed to measure statistical significance. For ROS measurements from acute brain slices *in vivo*, one-way ANOVA followed by Dunnett’s multiple comparisons test was performed for statistics. For mass spectrometry two-way ANOVA with Tukey’s post-hoc analysis was performed to measure significance.

## Results

### Increased ROS in astrocytes of a mouse model of AD

Previous studies have demonstrated an increase in ROS in the brains of both humans with AD and mouse models of AD (Testa et al., 2016; Yutuc et al., 2020). Increased oxidative stress in astrocytes has been implicated in the pathogenesis of AD (González-Reyes et al., 2017). To better understand this process, we applied a genetically encoded ratiometric hydrogen peroxide sensor, HyPerGR_T_, to examine hydrogen peroxide levels in the cytoplasm of astrocytes. In the presence of oxidation, HyPerGR_T_ exhibits an increase in green fluorescence, while maintaining relatively stable red fluorescence. This characteristic enables ratiometric calibration, which takes into account the different expression levels of the sensor in individual cells. Our previous study validated the response of this sensor to hydrogen peroxide and generated AAVs with an hSyn promoter and serotype 9 for imaging mitochondrial hydrogen peroxide in neurons (Li et al., 2022). To enable the expression of the sensor in a different cell type, we generated a new AAV plasmid containing a truncated GFAP promoter (Liu et al., 2008) (GfaABC_1_D) to direct astrocyte-specific expression (**Fig 1a**). Next, we utilized the newly developed plasmid along with packaging plasmids to generate AAVs with serotype 5.

**Fig 1.**
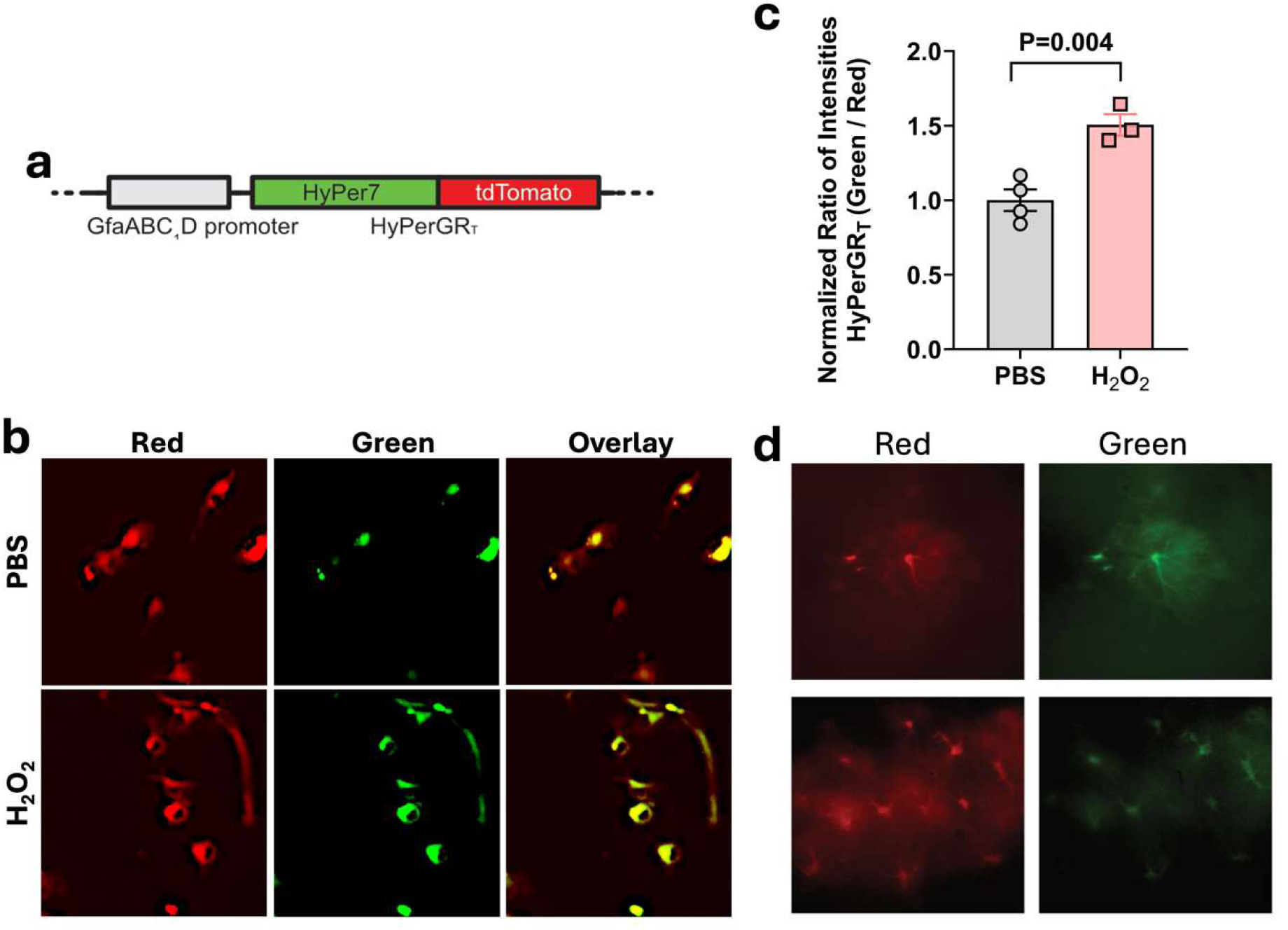
Astrocyte-specific expression of the HyPerGR_T_ sensor. (**a**) Arrangement of the genetic components for astrocyte-specific expression of HyPerGR_T_ driven by the GfaABC1D promoter. (**b, c**) Primary mouse astrocyte cultures expressing the AAV cytoplasmic HyPerGR_T_ sensor show increased ROS with hydrogen peroxide treatment. (**d**) Representative wide-field fluorescence images showing astrocyte-specific expression of the HyPerGR_T_ sensor in the brain of mice. Two different views are presented.

To validate the virus, we transduced primary astrocyte-enriched mixed glial cultures with the virus. After one week of viral expression, cells were treated with 100 mM hydrogen peroxide for 5 mins. Strong expression of the virus was observed in the cultured astrocytes (**Fig 1b**) and the green-to-red fluorescence ratio increased by approximately 50% following hydrogen peroxide treatment (**Fig 1c**). Next, to test the virus *in vivo*, we performed intracranial injections of the virus into the hippocampus of C57BL/6 mice. After 2-3 weeks of viral expression, strong fluorescence was observed in the acute brain slices obtained from these animals. The cells expressing the sensor exhibited morphological characteristics of astrocytes (**Fig 1d**). These results provide support for the robust astrocyte-specific expression achieved by AAVs carrying the GFAP promoter, which is consistent with previous findings (Griffin et al., 2019).

### Increased 7-ketocholesterol and ROS in mouse models of AD

Using our newly designed virus we determined the degree of oxidative stress in astrocytes in the AD brain. We conducted a comparison of the green-to-red fluorescence ratios between age-matched 5xFAD mice and control wild-type mice. The 5xFAD mice are known to have significant neuroinflammation (Forner et al., 2021). Following virus injection, 5xFAD mice displayed an increase in fluorescence ratios, indicating elevated ROS in astrocytes in these animals (**Fig 2a**).

**Fig 2.**
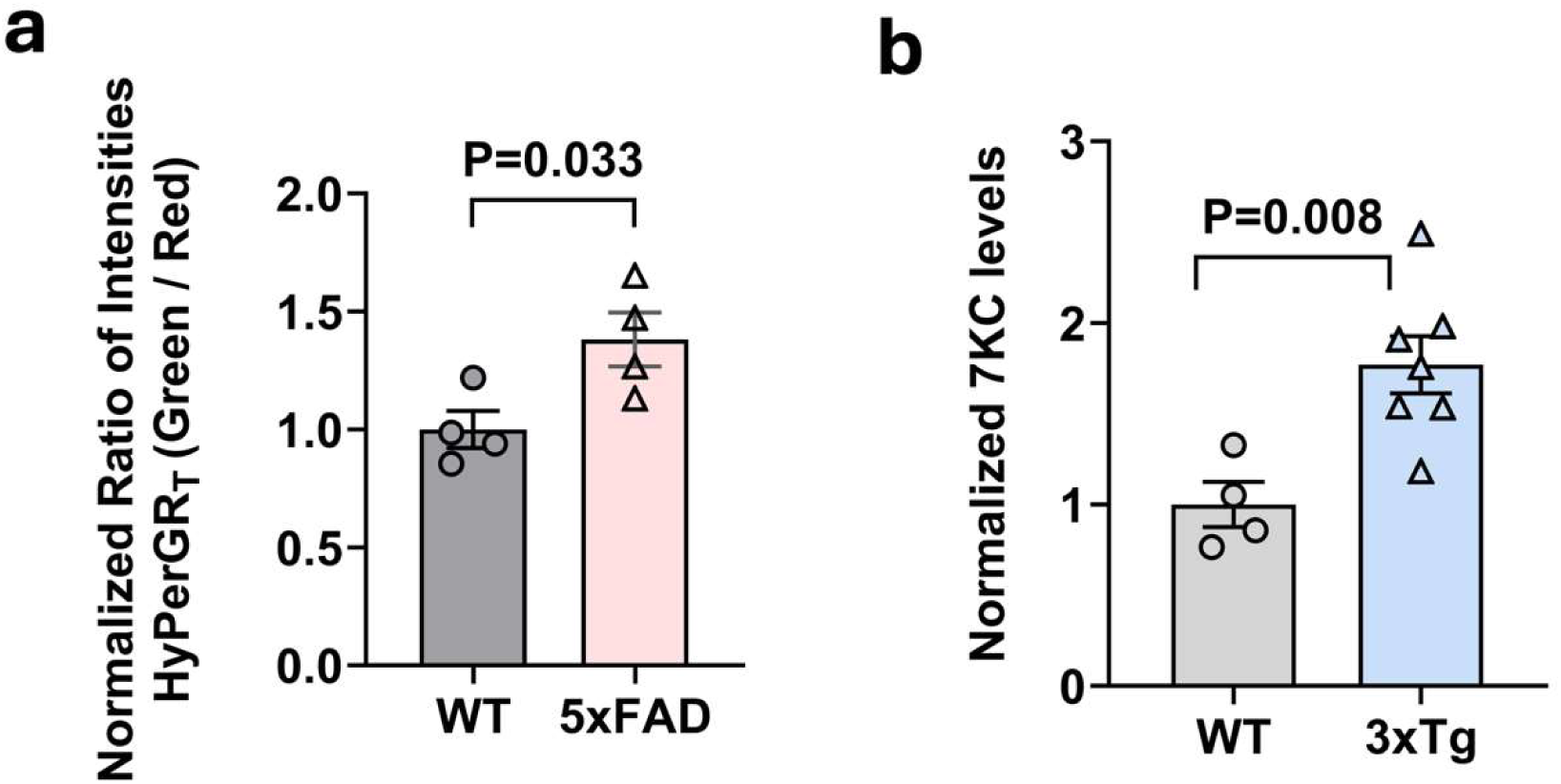
Increased ROS in Alzheimer’s disease mouse models. (**a**) Comparison of HyPerGR_T_ fluorescence (green-to-red ratio) in the hippocampus of heterozygous 5xFAD mice (HET, n=4) versus the wild-type control mice (WT, n=4). (**b**) 7-ketocholesterol is increased in 3xTg brain. Hypothalamus was isolated from 32-week-old female 3xTg and age-matched control mice. 7-KC was measured by mass spectrometry. Level was normalized to starting tissue weight. N=4-7 animals per group. Two-tailed unpaired t-test.

Because 7-KC is a product of oxidative stress and previous studies have demonstrated elevated levels of this oxidized cholesterol species in postmortem brains of individuals with AD (Testa et al., 2016), we assessed whether this phenomenon is also present in a mouse model of AD. We collected brains from 3xTg AD model mice and control B6129SF2/J mice. 3xTg mice express mutated forms of both human Aβ and tau (Oddo et al., 2003) (Gloria et al., 2021) and show evidence of neuroinflammation, but with slower progression than the 5xFAD model. Lipid samples were prepared from these brains and analyzed using mass spectrometry to determine the levels of 7-KC. The concentration of 7-KC was higher in the brains of 3xTg mice compared to controls (**Fig 2b**). Collectively, these findings suggest an association between Alzheimer’s pathology, oxidative stress in astrocytes and 7-KC.

### 7-ketocholesterol increases astrocyte ROS *in vivo*

Previous studies have implicated the oxidized cholesterol species 7-KC in driving oxidative stress in various diseases (Indaram et al., 2015). To investigate whether 7-KC can independently induce ROS in astrocytes *in vivo*, we conducted an experiment with C57BL/6J mice. We administered the astrocyte-specific AAV HyPerGR_T_ virus to the hippocampus of these mice. Following a 3-week period for viral expression, we performed stereotactic injections of 5 μM 7-KC or cholesterol (as a control) into the hippocampus. After 3 days, we imaged acute brain slices and observed a notable elevation of cytosolic ROS in astrocytes from the brains of mice injected with 7-KC, compared to those injected with cholesterol (**Fig 3**). This suggests that 7-KC can trigger an inflammatory response in the cytosol of astrocytes *in vivo*.

**Fig 3.**
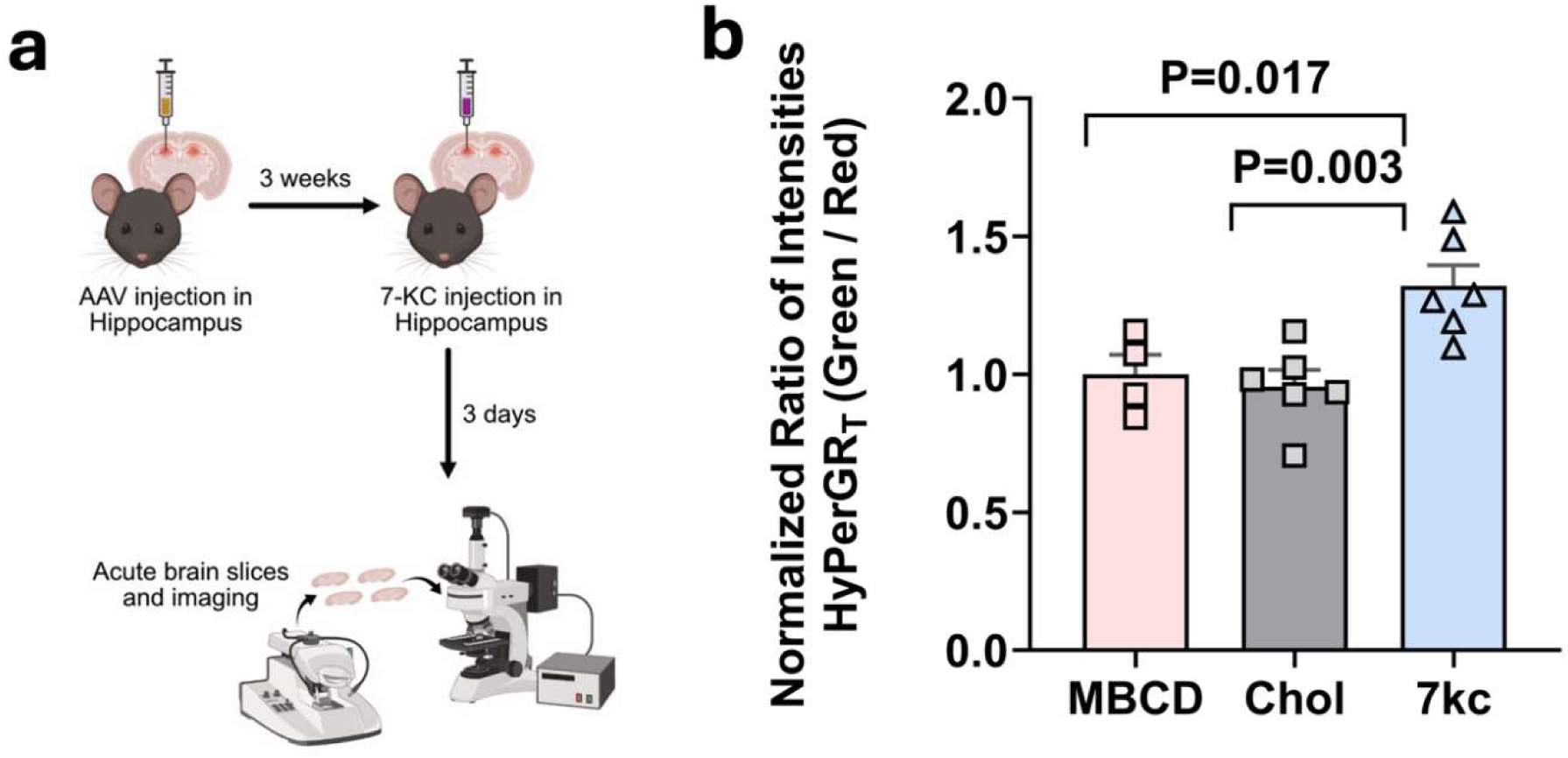
Imaging of 7-KC induced hydrogen peroxide in mice using the HyPerGR_T_ sensor. (**a**) Workflow of the experiment. 40-week-old C57BL/6 mice were injected with the AAV to express HyPerGR_T_ in hippocampal astrocytes. Three weeks later, 7-KC, cholesterol, or MBCD was injected into the brains of mice. In these experiments, MBCD was used as the vehicle for solubilizing 7-KC and cholesterol. Acute brain slices were prepared and imaged three days later. (**b**) Comparison of HyPerGR_T_ fluorescence (normalized green-to-red ratio) in the hippocampus of 7-KC treated mice (n=6), cholesterol-treated mice (n=6), and MBCD-treated mice (n=4). *P*-values were derived from one-way ANOVA followed by Dunnett’s multiple comparisons test.

### 7-ketocholesterol increases astrocyte ROS and activates astrocytes in a mixed glial culture system

To better understand the source of increased ROS in astrocytes we switched to an *in vitro* culture system. Astrocyte-enriched primary mixed glial cultures were transduced with astrocyte-specific AAV HyPerGR_T_ virus. After one week to allow for virus expression, cells were treated with 10 uM 7-KC or cholesterol for 24 hours and then imaged or collected for qPCR (**Fig 4a**). ROS was quantified using the green to red fluorescence ratiometry. We observed that astrocytes treated with 7-KC showed significantly higher ROS compared to cholesterol treated cells (**Fig 4b,c**). In addition to this, we performed qPCR for cytokines and activation markers of astrocytes and microglia. Transcripts of the widely used astrocyte activation marker, GFAP, were significantly increased with 7-KC, while GCLM and IL33 did not show significant increases (**Fig 4d**). We also observed that transcripts of the microglial gene TNFα, was significantly increased with 7-KC. IL1β and HMOX showed a trend towards increased expression in 7-KC treated cells (**Fig 4e**). These data indicated that 7-KC could be acting either directly on astrocytes or through microglia to increase astrocyte ROS.

**Fig 4.**
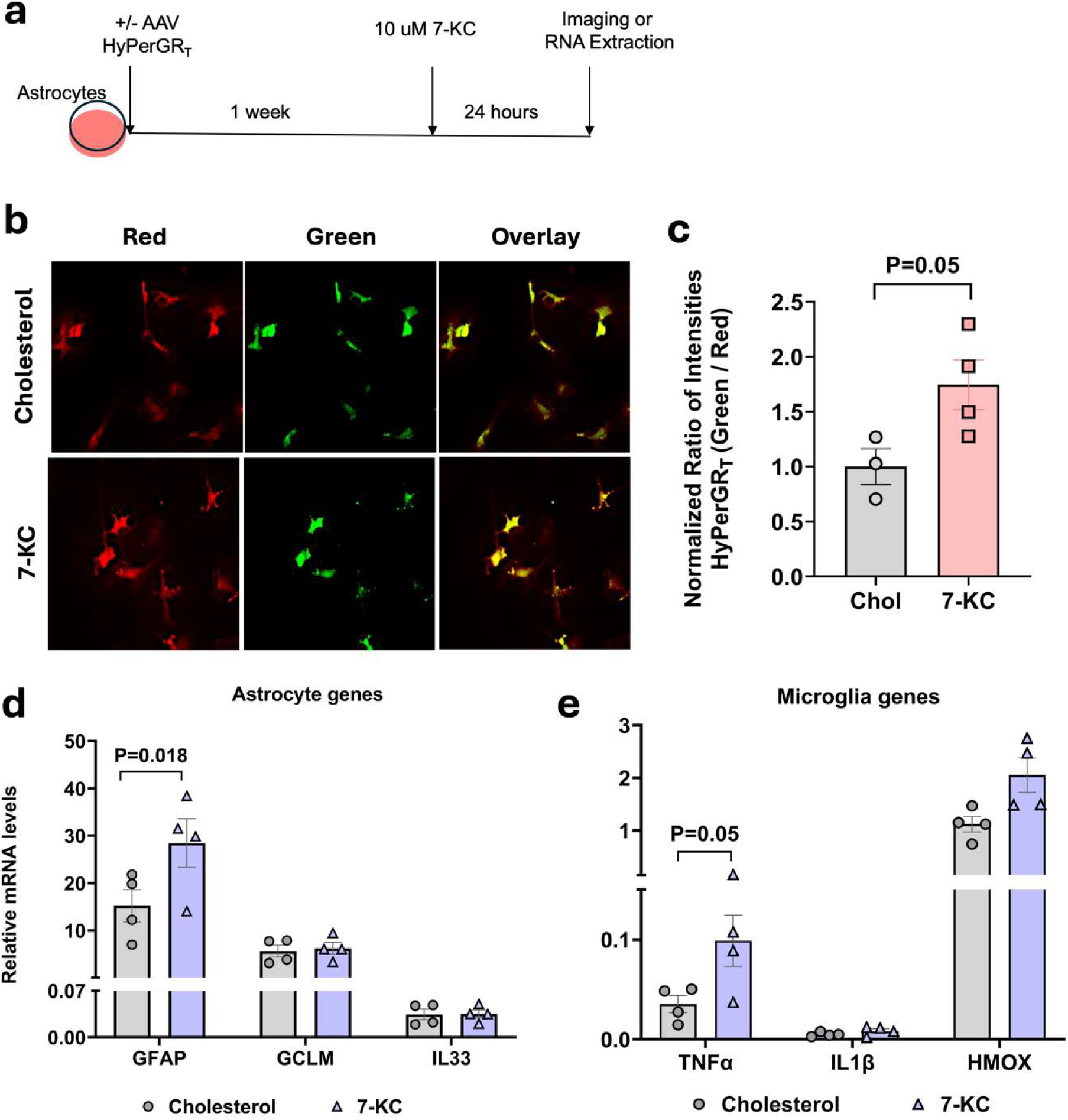
In vitro activation of astrocytes with 7-KC. (**a**) Experimental design showing 7-KC or cholesterol treatment on astrocytes (**b**) Astrocytes in mixed glial cultures expressing AAV HyPerGR_T_ treated with either 7-KC or cholesterol control. (**c**) 7-KC significantly increased astrocyte ROS as quantified by ratiometric fluorescence. (**d**) qPCR of astrocyte genes showing significant increase in GFAP with 7-KC treatment. (**e**) qPCR of microglia genes showing significant increase in TNFα with 7-KC treatment. All P values were obtained from two-tailed unpaired t-test.

### 7-ketocholesterol does not directly increase astrocyte activation markers

We next assessed if the increase in astrocyte activation observed in response to 7-KC was due to a direct effect of 7-KC on astrocytes. For this we treated primary mixed glial cultures with 5 μM PLX-3397 to significantly reduce microglial populations from the culture. After five days, the cells were treated with 7-KC or cholesterol for 24 hours and qPCR was performed (**Fig 5a**). In contrast to the mixed glial cultures, GFAP was not increased in response to 7-KC in the pure astrocyte cultures (**Fig 5b**). Transcripts of microglial markers TNFα, IL1β and HMOX were all very low and didn’t change between 7-KC and cholesterol treatments (**Fig 5c**), consistent with elimination of microglia from the cultures. These results indicates that 7-KC does not influence astrocytes directly and that microglia have a key role to play in the astrocyte activation by 7-KC.

**Fig 5.**
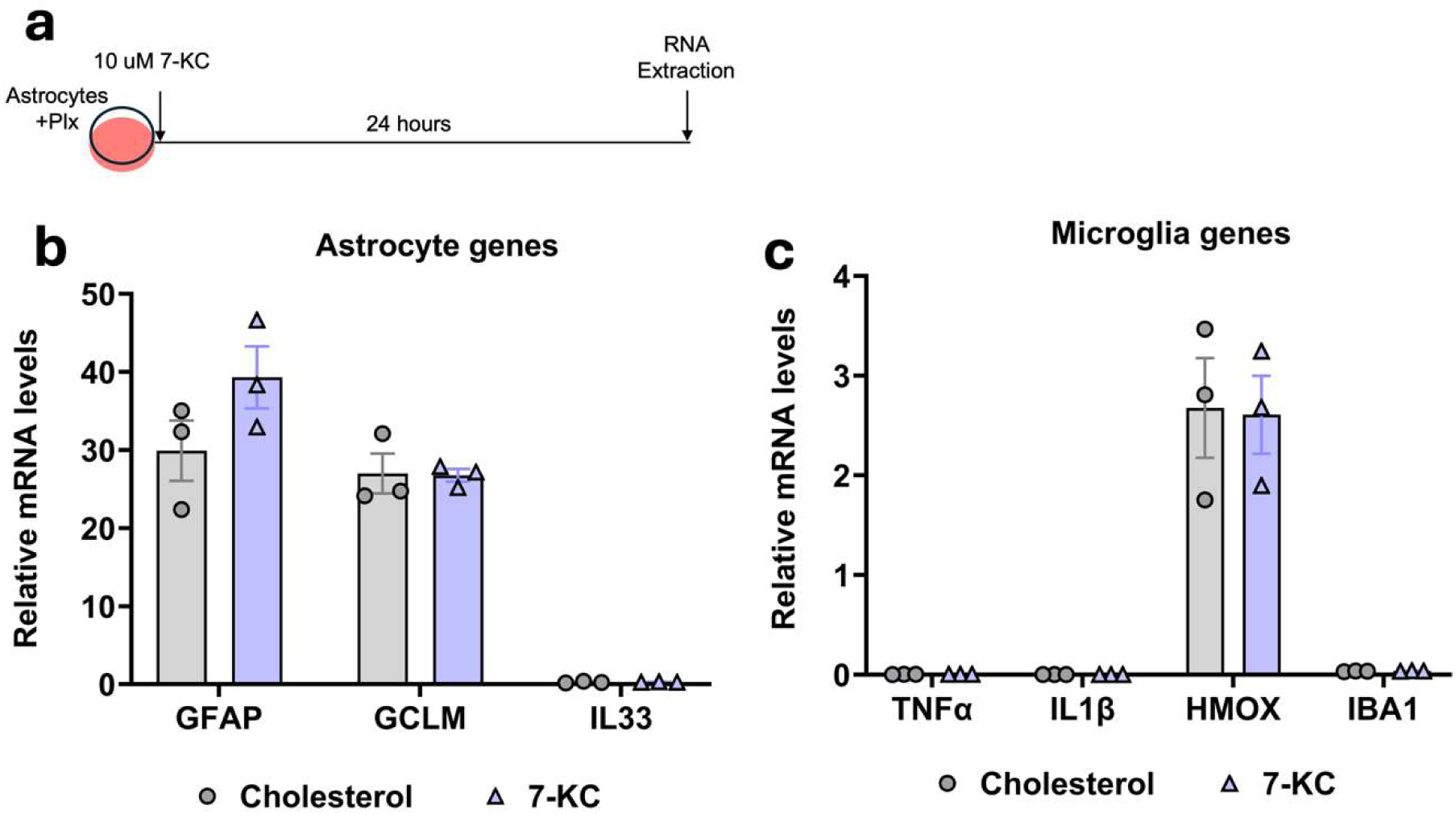
Microglia are required for astrocyte activation by 7-KC. (**a**) Experimental design showing 5μM Plx treatment and 10 μM 7-KC treatment in astrocytes. (**b**) qPCR of astrocyte activation markers with 7-KC or cholesterol control. (**c**) qPCR of microglial markers with 7-KC or cholesterol control. N=3 independent cultures.

### 7-ketocholesterol increases microglial activation markers in SIM-A9 cells

Given that we did not see significant activation of astrocytes with 7-KC when microglia were depleted from primary cultures, we investigated the activation of microglia directly with 7-KC. To test this, we primed SIM-A9 microglia-derived cell line with 0.5 μg/ml LPS, then treated with 12 μM of either 7-KC or cholesterol as a control (**Fig 6a**). In response to 7-KC, transcripts of microglia activation markers, IL1β and IBA1 were significantly increased (**Fig 6b**). TNFα transcript showed a trend towards increased expression, but did not reach statistical significance. Therefore, our results demonstrate that 7-KC is able to activate microglia directly.

**Fig 6.**
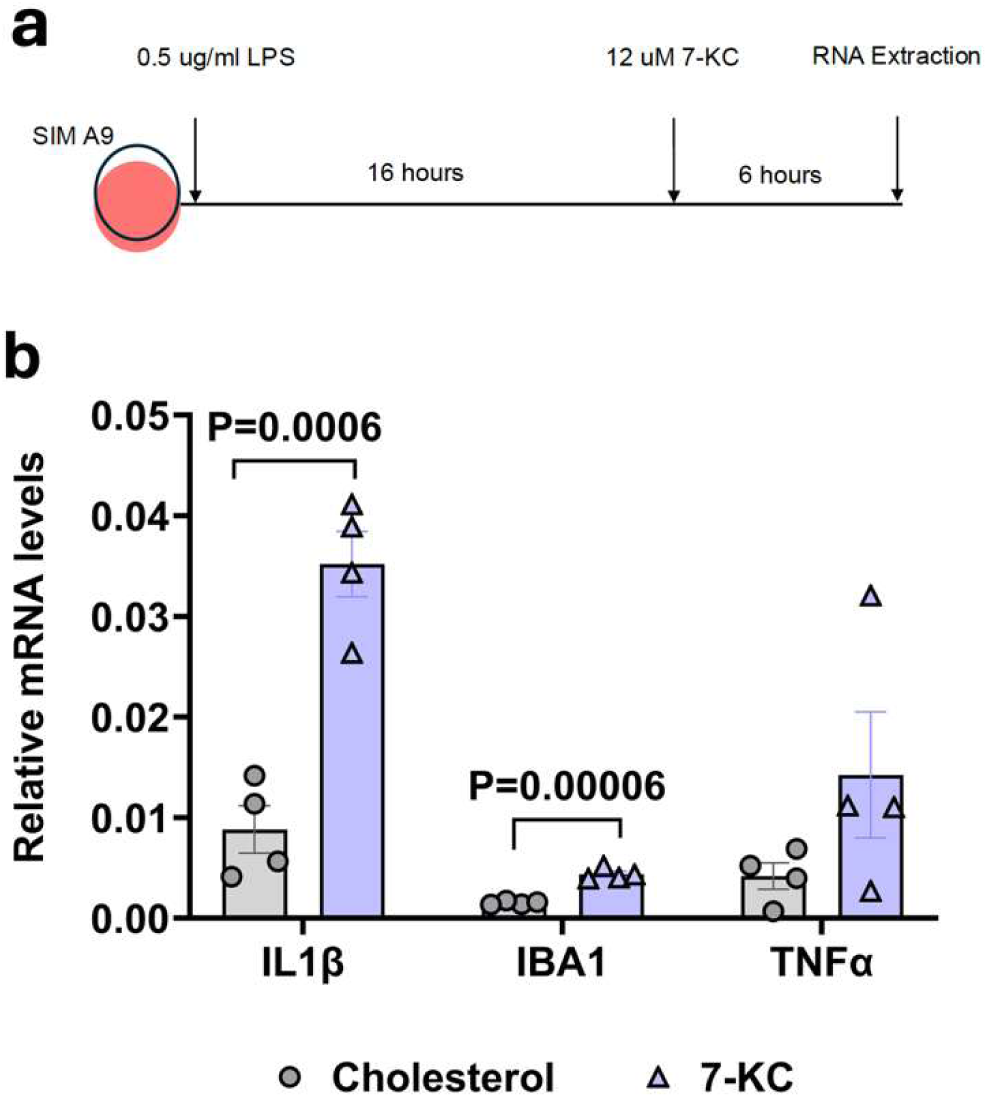
7-KC increases activation of SIM-A9 cells. (**a**) Experimental design showing 16-hour priming of SIM-A9 cells with 0.5 μg/ml LPS followed by 6-hour treatment with 12 μM 7-KC or control cholesterol (**b**) qPCR showing significant increase in IL1β and IBA1 transcripts with 7-KC. P-values were obtained from two-tailed unpaired t-test.

### Depletion of microglia from 3xTg mice is accompanied by reduced 7-KC

Microglia are the immune cells of the CNS and are activated in AD. Because 7-KC is generated in the presence of oxidative stress, we hypothesized that microglia would be a major source of 7-KC in AD. In order to investigate if increased 7-KC in 3xTg brains is initiated by microglia, we depleted microglia in these mice by feeding with PLX-3397. Following microglia depletion, brains were extracted and prepared lipid samples were analyzed for total cholesterol and 7-KC. As anticipated, 7-KC levels decreased in 3xTg brains upon microglia depletion, without impacting total cholesterol (**Fig 7a,b**). In conclusion, these data show that microglia are required for the elevations in 7-KC seen in AD and that this elevated 7-KC contributes to the increase in astrocyte ROS in this disease.

**Fig 7.**
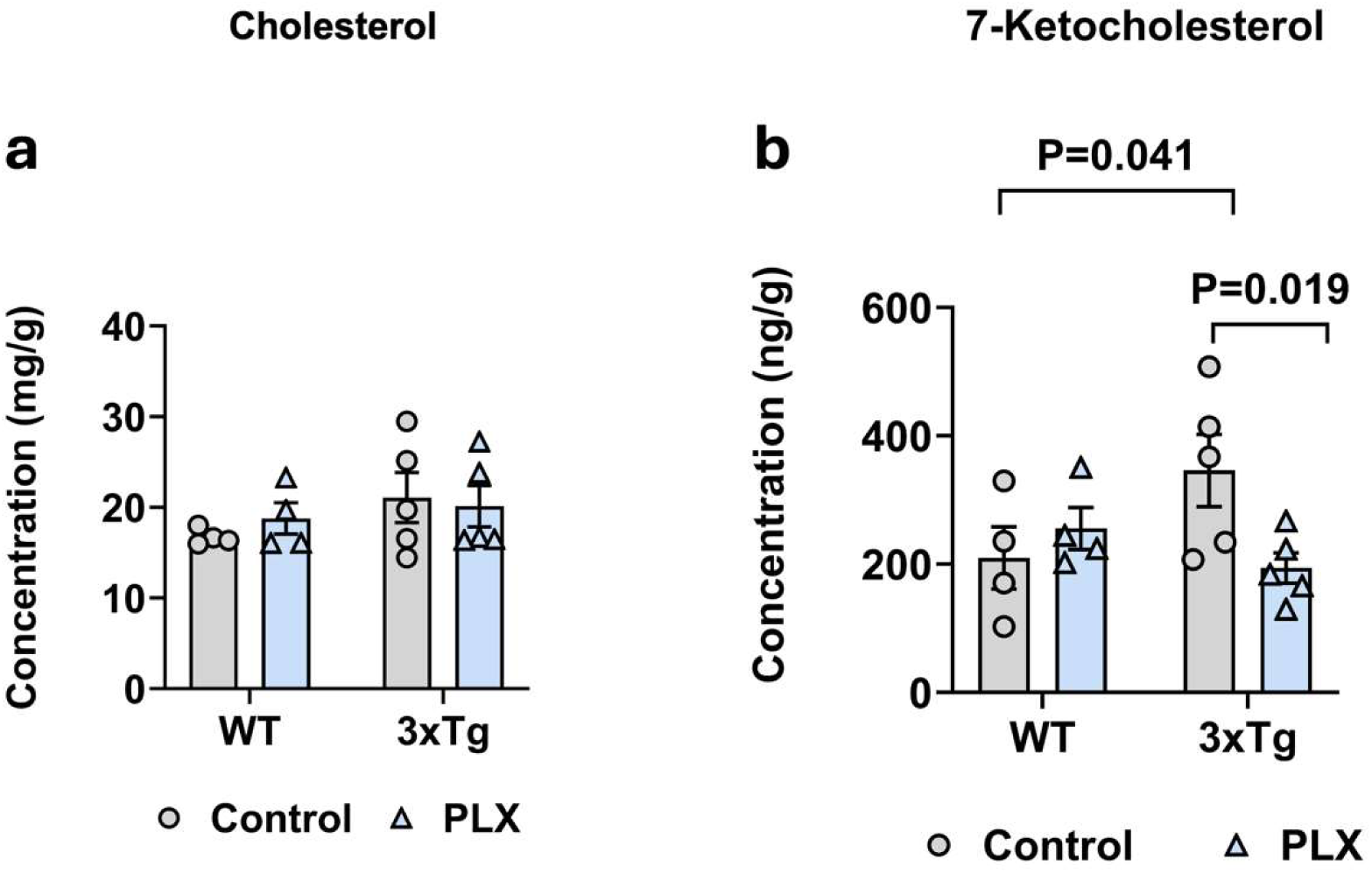
Microglia are required for the 7-KC increase in 3xTg brain. Hippocampi were isolated from 55-week-old female 3xTg and control mice fed either a control diet or Plexxikon-3397 for 4 weeks. **(a)** There was no difference in total cholesterol between groups, **(b)** however, there was a reduction in 7-ketocholesterol in the Plexxikon-fed 3xTg mice compared to chow-fed 3xTG mice. N=4-5 animals per group.

## Discussion

### 7-Ketocholesterol increases in Alzheimer’s Disease

7-KC is a toxic derivative of cholesterol mainly produced by auto-oxidation during adverse physiological changes and pathological conditions (Kim et al., 2001; Vejux et al., 2020). 7-KC is elevated in brain tissues (Testa et al., 2016), cerebrospinal fluids and plasma of patients with neurological disorders such as Alzheimer’s disease (Mahalakshmi et al., 2021) and it is a biomarker of oxidative stress (Iuliano et al., 2003; Samadi et al., 2019; Seet et al., 2010). While 7-KC is produced by the interaction of reactive oxygen species with cholesterol, it also furthers the stimulation of ROS production. Since inflammation is a prominent feature of AD, we examined the levels of 7-KC in 3xTg mouse brains. As determined by mass spectrometry, the levels of 7-KC were higher in 3xTg compared to control mouse brains. The magnitude of this increase is similar to what is seen in human AD brains (Testa et al 2016).

### Microglia drive 7-ketocholesterol-mediated increase in astrocyte ROS

There are several studies examining the impact of 7-KC on microglia, however the impact on astrocytes had not previously been examined. In this study we showed, using the novel AAV cytoplasmic GFAP HyPerGR_T_ ROS sensor, that astrocyte ROS increases in the hippocampus of C57BL/6J mice upon 7-KC injection. This suggested that 7-KC by itself can cause activation of astrocytes, however, treatment of astrocytes *in vitro* with 7-KC had minimal effect. In contrast, the microglial cell line SIM-A9 and mixed glial cultures responded to 7-KC, implicating microglia in 7-KC-mediated increases in astrocyte ROS. Our data support recent findings that chronic activation of microglia by oxysterols, including 7-KC, is linked to the ongoing neuroinflammation seen in AD (Krishnan et al., 2024). This inflammatory response exacerbates neuronal damage, contributing to AD pathology.

This was further confirmed by *in vivo* depletion of microglia in 3xTg mice, where 7-KC levels decreased with the depletion of microglia. This also indicates that microglia could be involved in the production of 7-KC and contribute to a vicious activation cycle, where microglia induce 7-KC production, which in turn activates microglia and causes subsequent inflammatory responses in astrocytes. This study thus provides an important role of 7-KC as a contributor to microglial activation, leading to astrocyte ROS, which could play a crucial role in the cumulative effects of inflammatory processes ultimately leading to neurodegeneration in AD. Further investigations are needed to study the mechanism of activation of microglia by 7-KC and the mechanism by which 7-KC-stimulated-microglia cause astrocyte activation that add up to disease progression. We envision that integrating the fluorescent biosensor with AD mouse models would allow for longitudinal two-photon imaging of ROS levels through cranial windows in AD mice. This *in vivo* model could be utilized to further study the mechanisms involved in the interaction between microglia, 7-KC, and astrocyte activation, providing valuable insights into the progression of AD and potential therapeutic targets.

### 7-KC as a potential therapeutic target for reducing ROS in AD

Microglia are critical players in the brain’s innate immune system and play a key role in defending against AD pathology. In AD, chronic activation of microglia can lead to the release of pro-inflammatory cytokines and reactive oxygen species. While acute inflammation can help remove damaged cells, chronic neuroinflammation contributes to neurodegeneration by harming neurons (Chen & Colonna, 2022).

Microglia and astrocytes communicate with each other to maintain CNS homeostasis and respond to challenges. This communication occurs through various pathways, including direct cell-cell contact and secretion of signaling molecules, such as cytokines. Activated microglia release a variety of signaling molecules that activate astrocytes, leading to changes in neuroinflammation. In this study we have shown the direct and crucial role played by 7-KC on microglial activation following elevation of ROS in astrocytes. Thus, 7-KC may be an important contributor to propagation of the neuroinflammation seen in AD. This study opens a new avenue of research that calls for further investigation towards means of attenuating 7-KC as a therapeutic target in reducing ROS in AD.

